# Unmasking cellular response of a bloom-forming alga to virus infection by resolving expression profiling at a single-cell level

**DOI:** 10.1101/186981

**Authors:** Shilo Rosenwasser, Miguel J. Frada, David Pilzer, Ron Rotkopf, Assaf Vardi

## Abstract

Marine viruses are major evolutionary and biogeochemical drivers of microbial life in the ocean. Host response to viral infection typically includes virus-induced rewiring of metabolic network to supply essential building blocks for viral assembly, as opposed to activation of anti-viral host defense. Nevertheless, there is a major bottleneck to accurately discern between viral hijacking strategies and host defense responses when averaging bulk population response. Here we use *Emiliania huxleyi*, a bloom-forming alga and its specific virus (EhV), as one of the most ecologically important host-virus model system in the ocean. Using automatic microfluidic setup to capture individual algal cells, we quantified host and virus gene expression on a single-cell resolution during the course of infection. We revealed high heterogeneity in viral gene expression among individual cells. Simultaneous measurements of expression profiles of host and virus genes at a single-cell level allowed mapping of infected cells into newly defined infection states and uncover a yet unrecognized early phase in host response that occurs prior to viral expression. Intriguingly, resistant cells emerged during viral infection, showed unique expression profiles of metabolic genes which can provide the basis for discerning between viral resistant and sensitive cells within heterogeneous populations in the marine environment. We propose that resolving host-virus arms race at a single-cell level will provide important mechanistic insights into viral life cycles and will uncover host defense strategies.

## Introduction

Marine viruses are recognized as major ecological and evolutionary drivers and have immense impact on the community structure and the flow of nutrients through marine microbial food webs [1-5]. The cosmopolitan coccolithophore *Emiliania huxleyi* (Prymnesiophyceae, Haptophyta) is a widespread unicellular eukaryotic alga, responsible for large oceanic blooms [6, 7]. Its intricate calcite exoskeleton accounts for ~1/3 of the total marine CaCO3 production [8]. *E. huxleyi* is also a key producer of dimethyl sulfide [9], a bioactive gas with a significant climate-regulating role that seemingly enhances cloud formation [10]. Therefore, the fate of these blooms may have a critical impact on carbon and sulfur biogeochemical cycles. *E. huxleyi* spring blooms are frequently terminated as a consequence of infection by a specific large dsDNA virus (*E. huxleyi* virus, EhV) [11, 12]. The availability of genomic and transcriptomic data and a suite of host isolates with a range of susceptibilities to various EhV strains, makes the *E. huxleyi*-EhV a trackable host-pathogen model system with important ecological significance [13-20]}.

Recent studies demonstrated that viruses significantly alter the cellular metabolism of their host either by rewiring of host-encoded metabolic networks, or by introducing virus-encoded auxiliary metabolic genes (vAMG) which convert the infected host cell into an alternate cellular entity (the virocell [21]) with novel metabolic capabilities [22-27]. A combined transcriptomic and metabolomic approach taken during *E. huxleyi*-EhV interaction revealed major and rapid transcriptome remodeling targeted towards *de novo* fatty acid synthesis [18] fueled by glycolytic fluxes, to support viral assembly and the high demand for viral internal lipid membranes [28, 29]. Lipidomic analysis of infected *E. huxleyi* host and purified EhV virions further revealed a large fraction of highly saturated triacylglycerols (TAGs) that accumulated uniquely within distinct lipid droplets as a result of virus-induced lipid remodeling [27]. The EhV genome encodes for a unique vAMG pathway for sphingolipid biosynthesis, never detected before in any other viral genome. Biochemical characterization of EhV-encoded serine palmitoyl-CoA transferase (SPT), a key enzyme in the sphingolipid biosynthetic pathway, revealed its unique substrate specificity which resulted in the production of virus-specific glycosphingolipids (vGSLs) composed of unusual hydroxylated C17 sphingoid-bases [30]. These viral-specific sphingolipids are essential for viral assembly and infectivity and can induce host programmed cell death (PCD) during the lytic phase of infection [14, 31]. Indeed, EhV can trigger hallmarks of PCD, including production of reactive oxygen species (ROS), induction of caspase activity, metacaspase expression, changes in ultrastructure features and compromised membrane integrity [32-34].

The high metabolic demand for building blocks required to support synthesis, replication and assembly of large viruses with high burst size as EhV [34-36], point to high dependence of viruses on their host metabolic state for optimal replication [21, 37]. Consequently, heterogeneity in host metabolic states as a result of complex interactions between nutrient availability and stress conditions may affect the infection dynamics. However, almost all of our current understanding of the molecular mechanisms that govern host-virus interactions in the ocean, is derived from experiments carried out at the population level, assuming synchrony and uniformity of the cell populations and neglecting any heterogeneity. Additionally, averaging the phenotypes of a whole population hinders the investigation of essential life cycle strategies to evade viral infection that can be induced only by rare subpopulations [38]. Understanding microbial interactions at a single-cell resolution is an emerging theme in microbiology. It enables the detection of complex heterogeneity within microbial populations and has been instrumental to identify novel strategies for acclimation to stress [39-41]. The recent advancement of sensitive technologies to detect gene expression from low input-RNA allows quantification of heterogeneity among cells by analyzing gene expression at the single cell level [42, 43]. High-throughput profiling of single-cell gene expression patterns in mammalians and plant cells led to the discovery of new cell types, detection of rare cell subtypes, and provides better definition and cataloging of developmental phases in high resolution [44-48]. Importantly, the role of cell-to-cell communication and variability in controlling infection outcomes has only been recently demonstrated in cells of the mammalian immune system in response bacterial pathogens [49-52]. Cell-to-cell variability in host response to viral infection was observed in several mammalian viruses and was attributed to several factors, including intrinsic noise (e.g. stochasticity of biochemical interactions involved in the infection process), the number of viral genomes initiating the infection process and the specific cell-state before the infection [52-55].

Recently, simultaneous detection of host and pathogen gene expression profile was suggested as a powerful tool used to gain a better understanding of the molecular mechanisms underlying the infection process and to identify host resistance responses [21, 56-58]. However, the existence of cell-to-cell variability during infection suggest that key events in host response are masked by conventional bulk cell expression profiling and that detection of gene expression on single cell resolution may uncover hidden host responses.

Here, we quantified the dynamics of host and virus gene expression profiles of individual cells during infection of *E. huxleyi* populations. We provide strong evidence for heterogeneity within the population and discern between cells at different infection states based on their viral gene expression signatures. We unravel an unrecognized phase of early host response that preceded viral gene expression within infected cells. We suggest that examining host and virus gene expression profiles at the single cell resolution allows to infer the temporal dynamic of the infection process, thereby it serve as an attractive approach to decipher the molecular mechanism underlying host-virus interaction.

## Results and Discussion

To examine the variability within infected *E. huxleyi* cells, we measured the expression levels of selected host and viral genes over the course of infection at a single-cell resolution. Cells were isolated during infection of *E. huxleyi* CCMP2090 at different phases, at 0, 2, 4, 24 hours post infection (hpi) (Figure 1). We used the C1 single-cell Auto Prep System to sort and extract RNA from single *E. huxleyi* cells during viral infection by EhV201). The presence of a single cell captured in an individual isolation chamber was confirmed by microscopic inspection of emitted chlorophyll auto-fluorescence (Figure 2A). In order to detect variability in viral infection states, we conducted simultaneous measurements of expression profiles of host and virus genes at a single-cell level by using multiplexed qPCR. We selected viral genes encoding for sphingolipid biosynthesis as well as gene markers for early and late infection[18, 59]. Selected genes involved in host metabolic pathways were targeted based on previous reports which demonstrated their functional role during infection, including primary metabolism (glycolysis, fatty acid biosynthesis), sphingolipid and terpenoid metabolism, autophagy and antioxidant genes [18, 27, 33, 34]. In addition, we examined the expression of host genes associated with life cycle, meiosis and PCD [32] that exhibited induction during infection [60], (see Supplemental Table 1 for primers list).

**Figure 1:**
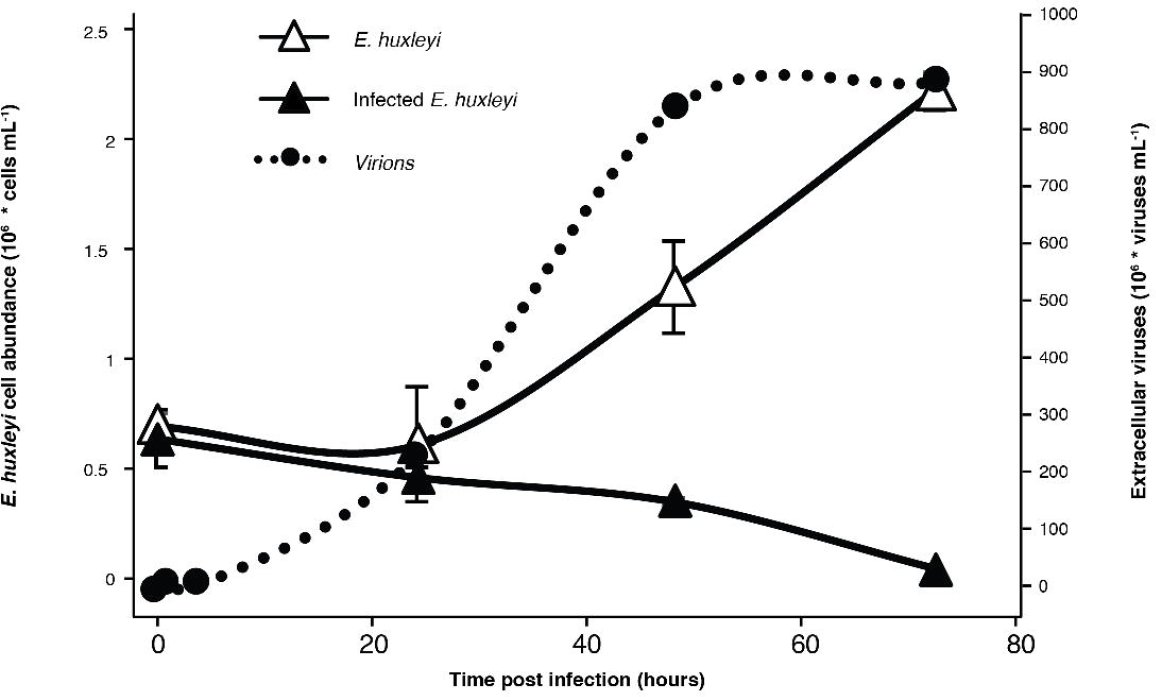
Infection dynamics of *E. huxleyi* by its specific virus EhV. *E. huxleyi* CCMP2090 culture was infected by the EhV201 lytic virus and compared with uninfected control cells. Host cell abundance and production of extracellular viruses were monitored using flow-cytometry. (mean ± SD, n = 3, at least 6000 cells were measured at each time point).

**Figure 2:**
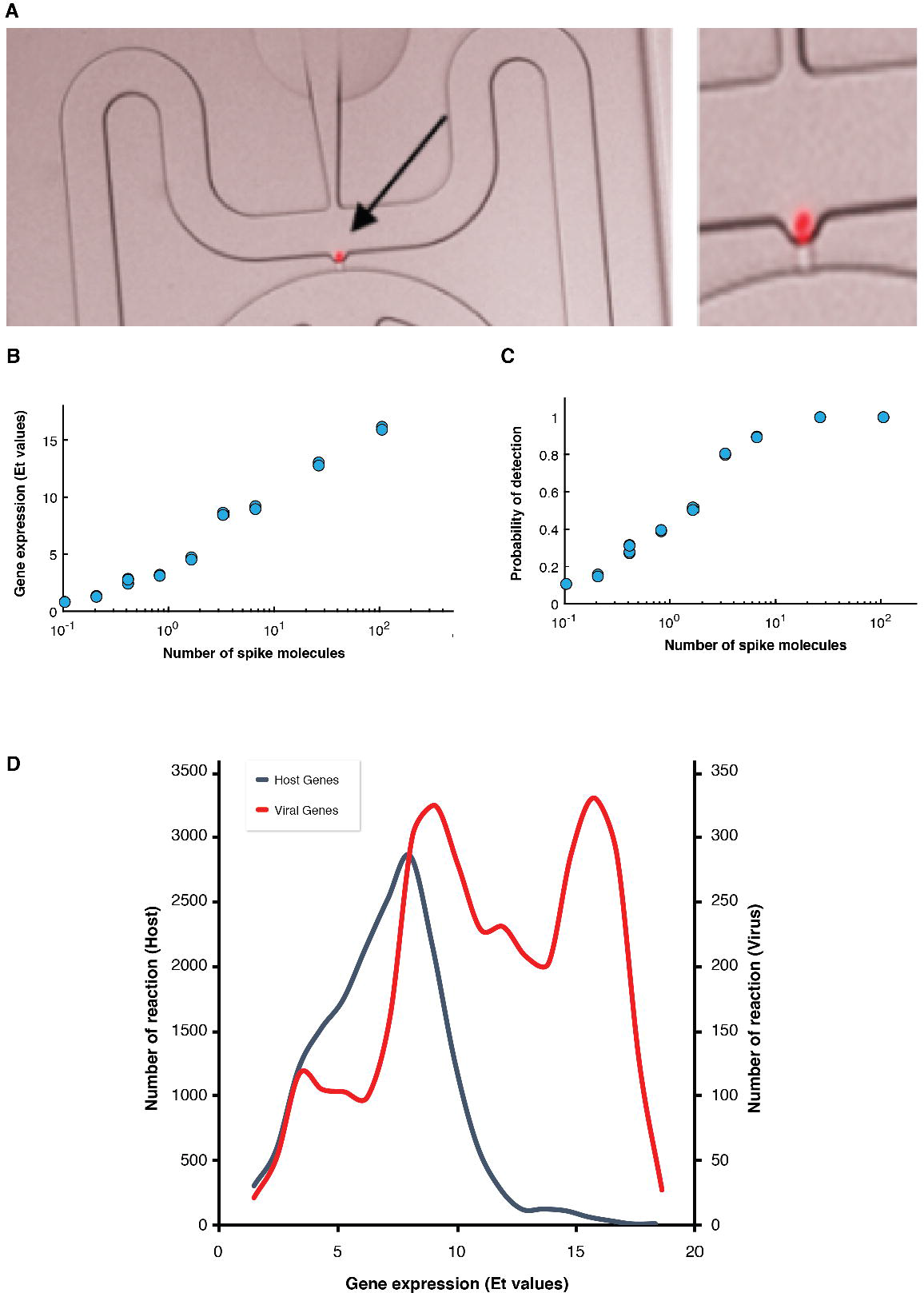
Host and virus gene expression profiling at a single cell level. (A) Automated microfluidic capture of a single *E. huxleyi* cell in the C1 chip (red: chlorophyll autoflouresence, indicated by a black arrow), the image on the right is a zoom into the image of a single cell. (B,C) Examination of detection level of single-cell gene expression analysis. A set of ERCC RNA molecules were spiked to each C1 well and their level was determined using multiplex qPCR. (B) The fraction of wells with positive qPCR reaction (Ct < 30) for each examined spike. (C) The correlation between the average level of expression (Et) value and the number of spike molecule. (D) Distribution of host and virus genes expression among individual cells. The average expression values of host and viral genes among isolated single cells was calculated and the distribution is presented.

To test for the sensitivity in detection of gene expression on a single cell level, we spiked-in, to each C1 well, a set of External RNA Controls Consortium (ERCC) molecules that span a wide range of RNA concentrations (from ~0.5 to ~100 molecules per well). We subsequently quantified their concentration using similar qPCR amplification setup as used for the host and virus genes. Pairwise correlation between spike concentrations and Et (Et=30-Ct) values obtained from the qPCR was >0.98 (Pearson correlation coefficient, p-value= 4.2^.10−12^, Figure 2B). We found a highly sensitive level of detection with 40% probability to detect an RNA spike that is at a level of 1 molecule per sample (Figure 2C), similar to the detection level reported for mammalian cells [61]. Mean expression of viral and host genes in all examined cells were found to be 11.8 ± 4.0 and 6.96 ± 2.5 (Et values ± SD), respectively (Figure 2D).

We detected a high variability in viral expression profiles among individual cells within the same infected population. For example, heterogeneity in the expression levels of virus-encoded ceramide synthase (*vCerS*, EPVG_00014), a key enzyme in sphingolipid biosynthesis [18, 30] was detected during early phase of infection (2 and 4 hpi of CCMP2090, Figure 3A). Similar results were obtained for the average expression of 10 viral genes (Figure 3B). At the onset of viral lytic phase (24 hpi), all of the examined cells showed high viral gene expression (Figure 3A&B), suggesting that viruses eventually infected all of the examined host cells. Nevertheless, we cannot exclude the existence of a rare subpopulation that did not express viral genes. Principal component analysis (PCA) of viral gene expression among individual host cells showed that infected cells are distributed across distinct states of viral expression levels (Figure 3C). All viral genes had positive and similar coefficients for the PC1 component which captures ~90% of the variability of viral gene expression and found to be highly correlated to the average viral infection level (r = 0.99, Pearson linear correlation). These results demonstrate that PC1 reflected the intensity of viral infection. Accordingly, we used the score value of PC1 as an index for the level of expression of viral genes in each individual cell and termed it “infection index”.

**Figure 3:**
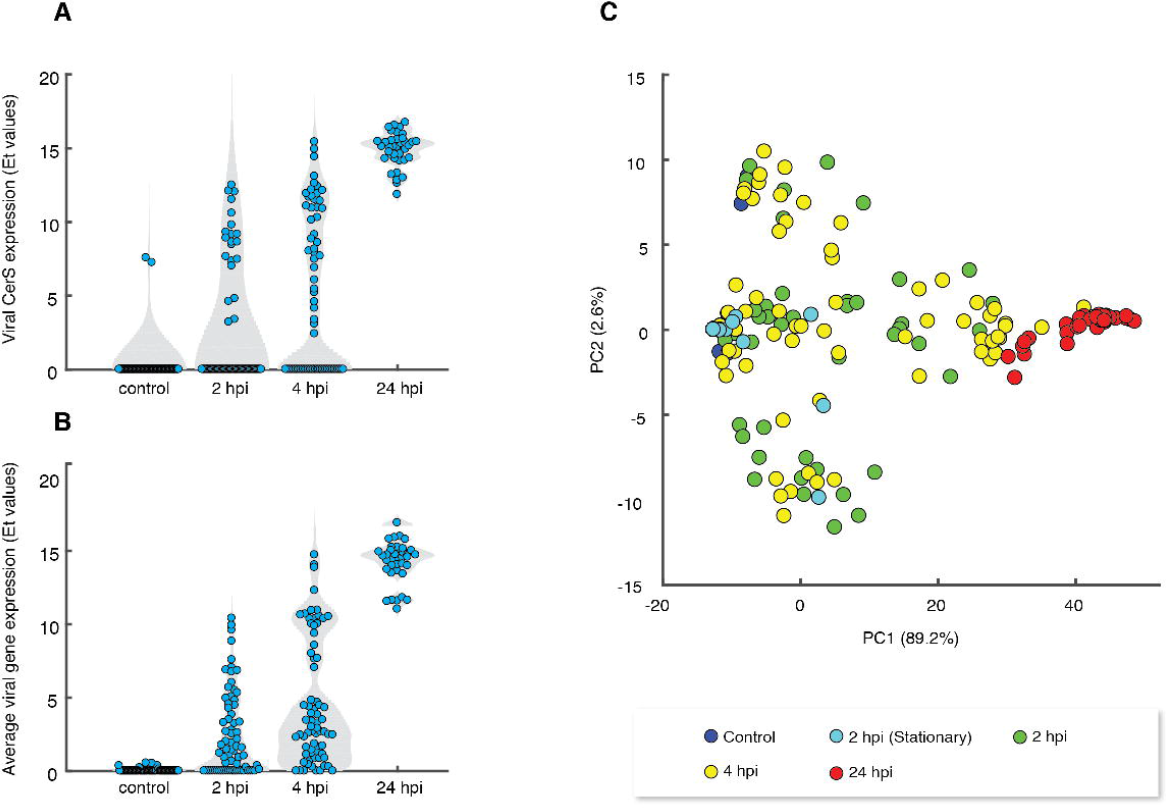
Single-cell analysis of infected population unmasks heterogeneity in viral gene expression profiles. (A) Violin plots of the expression value (Et) of viral dihydroceramide synthase (vCerS, EPVG_14, Gene bank: AET97902.1) at different hours post infection (hpi) of CCMP2090 cells infected by EhV201. (B) Violin plots of the mean expression value of 10 viral genes at different times post infection of CCMP2090 cells with EhV201. (C) Principal component analysis (PCA) plots of gene expression profiles of 10 viral genes derived from 323 individual E. huxleyi cells that were isolated from infected CCMP2090 cultures at different hpi.

We further realized that averaging host phenotypes over the course of infection might hinder our ability to observe the initial response of the host to viral infection and that single-cell analysis could significantly increase the resolution for sensitive detection of host response at this early stage of infection. We therefore re-ordered infected cells based on their viral infection index (PC1), rather than the actual time of infection (i.e. hpi), resulting in “pseudotemporal” hierarchy of single cells. Intriguingly, we unmasked a fraction of cells that were exposed to the virus but did not exhibit any detectable expression of viral genes. These cells had similar infection index values as control cells, with PC1 values <−10. We found that 33/179 (17%) of infected cells of CCMP2090 were at this distinct “lag phase” of viral infection. These individual cells were analyzed for their respective host gene expression levels based on a sliding window approach as it is less sensitive to technical noise which often observed in single cell data. We also used a statistical model to test for genes that are differentially expressed at these early stages of viral infection. This model incorporates the two types of heterogeneity that usually appear in single cell data, namely, the percentage of cells expressing a gene in a given population (e.g. Et value > 0) and the variability in expression levels in cells exhibiting positive expression values [62]. Up-regulation of several host genes in infected cells was detected prior to viral expression (Figure 4A-C and supplemental Table 3). An intriguing example is the *metacaspase-2* gene (*p*= 0.0000027) which was previously suggested to be induced and recruited during EhV lytic phase and activation of *E*. *huxleyi* PCD [32]. We also found early induction of triosephosphate isomerase (*TPI*, *p*=0.00063) and phospholipid:diacylglycerol acyltransferase (*PDAT*, *p* = 0.0018) which are involved in glycolysis and TAG biosynthesis. In addition, genes involve in autophagy [34] and *de novo* sphingolipid biosynthesis [18, 30] were detected in this unique early phase of host response. Since all of these metabolic pathways were recently shown to be essential for EhV infection[14, 18, 20, 21, 27, 30, 31, 33, 34], early induction of these pathways may serve as an effective viral strategy to prime optimal infection. Alternatively, this phase of early host response prior to viral gene expression may represent a newly unrecognized phase of immediate host anti-viral defense response. At the late stages of infection (infection index >10), we observed induction of several meiosis-related genes, including *HOP1* and *MND* and two *SPO11* homologues and *MYB* in CCMP2090 (Figure 4B). These results are in agreement with previous studies that suggested a phenotype switch of *E. huxleyi* to evade viral infection [38] and propose the induction of meiosis-related genes as part of transcriptomic reprogramming of during infection [60].

**Figure 4:**
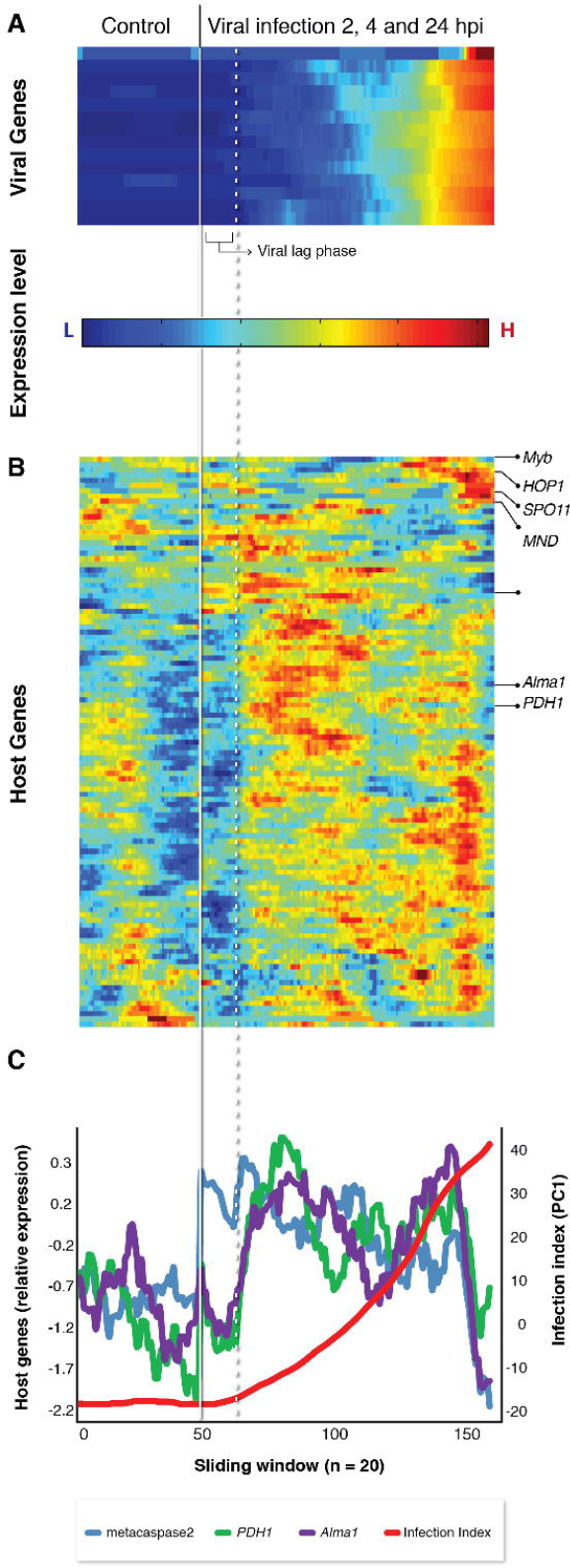
Host-virus co-expression patterns across viral infection states. Cells were re-ordered based on their infection index to reconstruct pseudotemporal separation of the infection process. (A, B) Clustogram representation of the average expression value of viral (A) and host (B) genes across the infection dynamics of CCMP2090 using a sliding window approach (window size = 20 cells). (C) Expression profile of selected host genes along the viral infection index (PC1) in the sliding windows of 20 cells reveals early induction of host genes prior to viral gene expression.

Further inspection of the PCA analysis showed the cells exhibiting low to moderate level of PC1 were highly variable in their PC2 level. Interestingly, we found a positive correlation (r = 0.53) between PC2 and the expression level of viral RNA polymerase gene (EPVG_00062) which was previously reported to be expressed at early-mid phases of infection [18, 59], while a negative correlation (r = -0.44) was found for a viral gene (EPVG_00010) that is known to be expressed at late phases of infection. Accordingly, cells with low PC2 levels expressed EPVG_00010 and not EPVG_00062, while cells with high PC2 values exhibited the opposite trend (Figure 5A and B).

**Figure 5:**
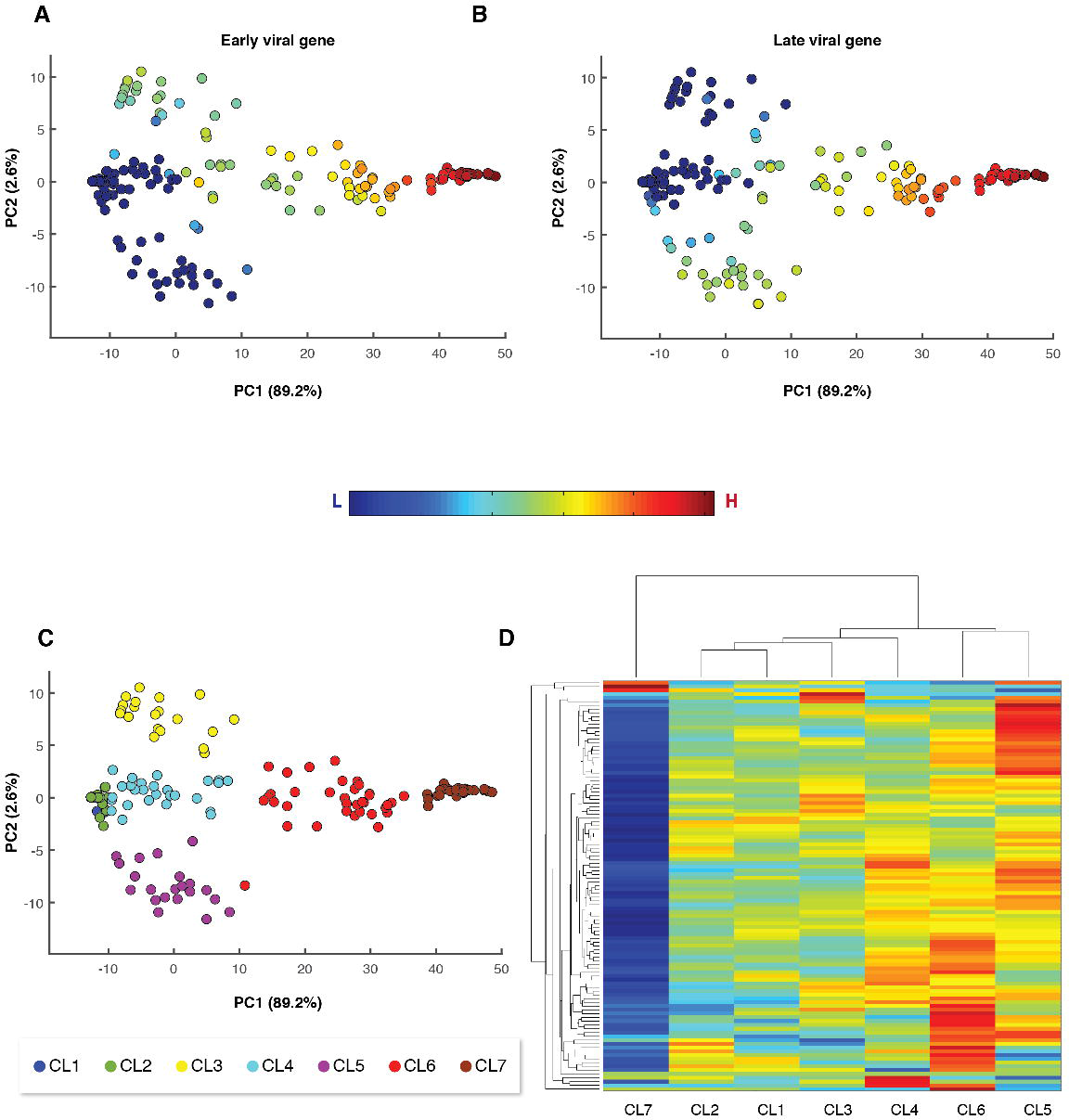
Viral expression is associated with induction of host metabolic genes at distinct phases of infection. (A, B) The same PCA plots as in Figure 3C with overlay, by a color code, representing the expression level of viral genes (Et values) that are associated with early-mid (A) and late (B) phases of viral infection (EPVG_00062 and EPVG_00010, respectively). (C) The same PCA plots as in Figure 3C with overlay, by a color code, representing the different clusters. Cells were clustered manually based on their infection index (PC1) and PC2 scores. (D) Clustogram representation of the of expression values of 109 host metabolic genes in the different clusters (defined in A). (D) Clustorgam representation of expression values of 109 host metabolic genes across the different clusters as defined in (C).

To further characterize host gene expression during different phases of infection, we manually clustered CCMP2090 cells according to their infection index (PC1) and the expression of either early or late viral genes (PC2) (Figure 5C) and examined the expression of host metabolic genes in these clusters (Figure 5D). This analysis showed that induction of most of host metabolic genes occurred in cells that expressed predominantly late viral genes (Figure 5D, CL5, -10<PC1<10, PC2>5) and in cells with moderate expression of viral genes (Figure 5D, CL6, 10<PC1<30). Down-regulation of many host genes was found in cells exhibiting high viral expression (Figure 5D, CL7, PC1>30), suggesting that these cells were at the final stages of infection.

In order to further characterize the link between optimal host metabolic state and efficient viral infection, we infected CCMP2090 stationary culture and subjected single cells to dual gene expression analysis at 2 hpi (Figure 6A). While most of the exponential growing cells exhibited viral expression, we detected only moderate viral expression in 3/27 (11%) of the stationary phase cells (Figure 6A), while the rest of the cells had viral expression patterns similar to uninfected cells (control). In parallel, stationary phase cells (either control or infected) exhibited down-regulation of most of the examined host metabolic genes, in contrast to their general up-regulation in infected exponential phase cells (Figure 6B). These data suggest that the cell-to-cell variability in host metabolic state may play important role in determining susceptibility to infection by large viruses with high metabolic demand. “Kill the Winner” is a key theory in microbial ecology which suggests that viruses shape diversity of microbial populations by infecting the most dominant proliferative host [63]. We propose that “Kill the Winner” may even act within isogenic populations based on the variability in the metabolic state, which will lead to differential susceptibility to viral infection, forming continuous host-virus co-existence [64]. It is possible that cell-to-cell heterogeneity in the metabolic activity is shaped by the tradeoff between complex abiotic stress conditions (e.g. nutrient availability [65-67] and light regime) and biotic interactions (e.g. pathogenicity or allelopathy), and may result in differential susceptibility to viral infection in the marine environment.

**Figure 6:**
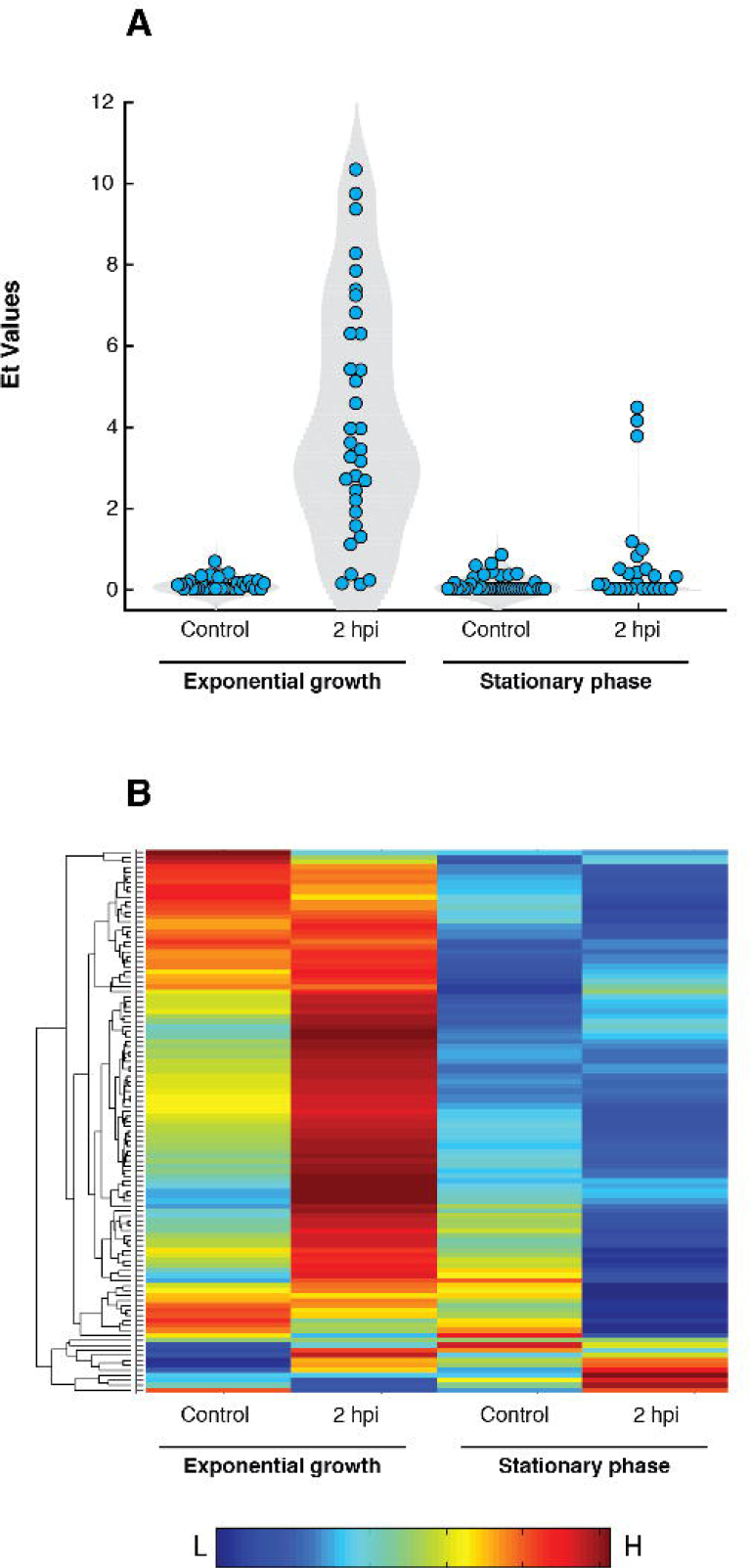
Low viral gene expression is culture at stationary phase is associated with dowb-regulation of host metabolic genes. (A) Violin plots of the mean expression of viral genes in individual exponential and stationary cells at 2 hpi and in uninfected cells (Control). (B) Clustorgam representation average expression values of 109 host metabolic genes in individual exponential and stationary cells at 2 hpi and in uninfected cells.

We further investigated whether uninfected sensitive and resistant *E. huxleyi* cells exhibited altered expression profiles in the host metabolic genes that showed variable expression during infection (Table S4). We exposed *E. huxleyi* cultures to viral infection and isolated cells that acquired resistance to subsequent viral infection of diverse EhV isolates (Figure 7A, [60]). We compared the expression profiles of recovered resistant cells (n = 18) to their mother cells that were highly susceptible to viral infection (n = 76). The tendency of resistance cells to aggregate make it difficult to isolate single cells, therefore for these analysis also doublet cells were included. Intriguingly, resistant and sensitive cells tend to cluster distinctively along the PC2 dimension (Figure 7B). Among the genes that drive the separation along the PC2 dimension and were differentially expressed in resistant and sensitive cells were *TPI*, diphosphomevalonate decarboxylase (*MVD1*) and *ceramidase-3* (Figure 7C) which are key enzymes in glycolysis, terpenoid and sphingolipid metabolism, respectively. Since *de novo* ceramide biosynthesis is uniquely encoded in the EhV genome, activation of ceramidase may serve as an anti-viral host response [18, 30]. Interestingly resistant cells also exhibited high expression of *metacaspase2* which was also highly expressed in cells with no viral expression in early phase of infection (Figure 7C). This data suggests that susceptibility to viral infection has a clear signature in expression profiles of host genes detected on a single-cell level.

**Figure 7:**
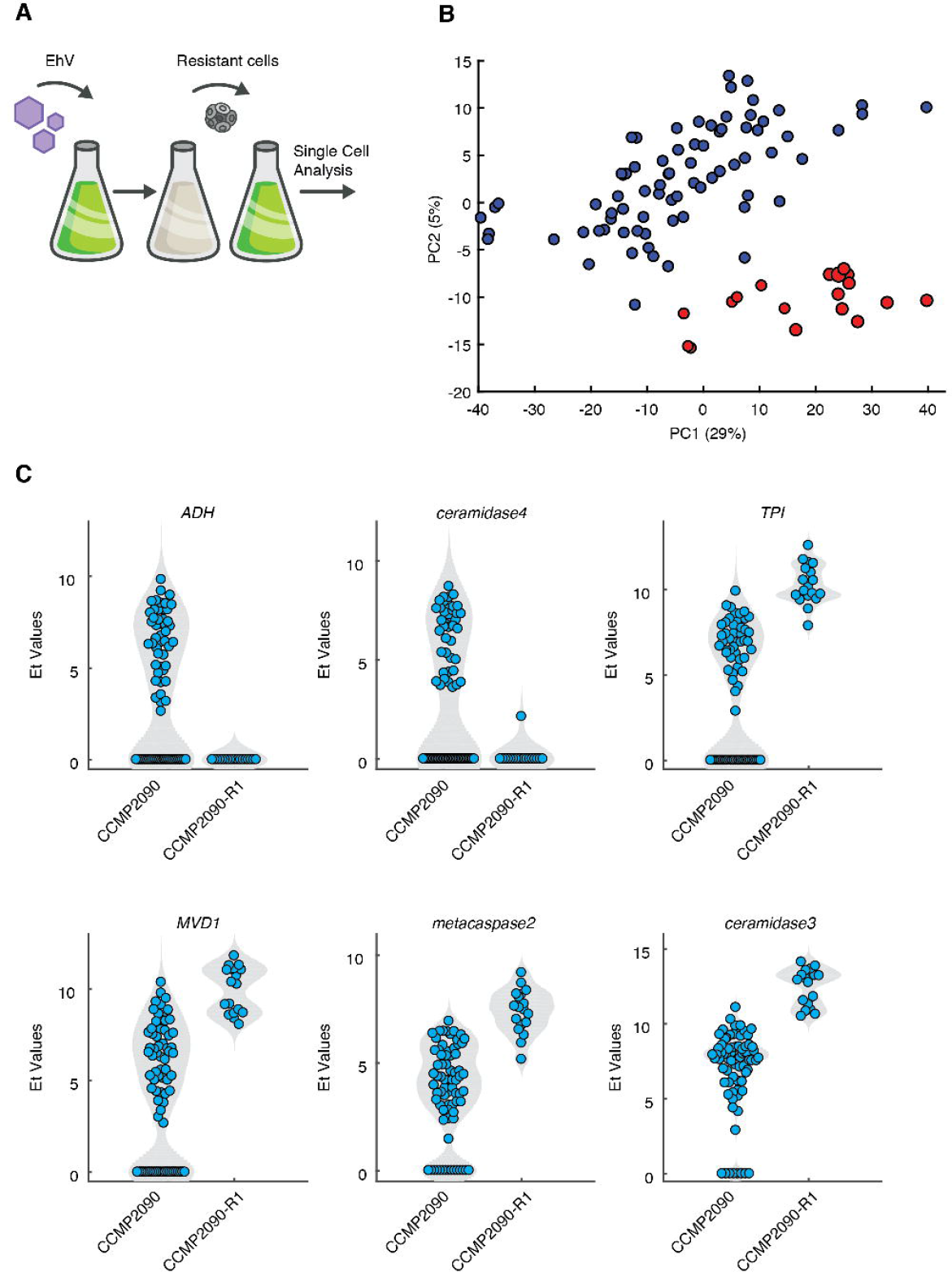
Differential expression of host gene on a single-cell level in virus-sensitive and virus-resistant cells. (A) Virus-resistant cells were isolated from infected CCMP2090 cells. (B) PCA projection of gene expression profiles of 93 host metabolic genes in from 94 individual *E. huxleyi* cells that were isolated from the sensitive CCMP2090 and resistance CCMP2090 culture (CCMP2090-R). Duplet cells are visualized by slightly bigger dots.(C) Violin plots of selected host genes that highly contributed to the separation of cells along PC2. The SingleCellAssay R package [62] was used to test significant changes in gene expression and all presented genes had *p-value* < 0.05 of hurdle test (See supplementary Table 4).

Although the mechanism for resistance of *E. huxleyi* to viral infection requires further investigations, the differential regulation of host metabolic genes suggests a unique specialized metabolism that differs from that of susceptible cells [68-70]. Future single-cell RNA-seq transcriptomic studies will provide high throughput identification of gene markers that are specific for resistant strains as well as new mechanistic insights into the molecular basis for resistance mechanisms.

## Conclusions

The data presented here suggests detection of host and virus expression profiles on a single-cell level as a novel approach to characterized host responses during viral infection in high resolution which is commonly masked in whole population RNAseq approaches[71]. By applying dual gene expression profiling during algal host-virus interactions, we uncovered an early host transcriptional responses. This newly defined phase can result from either induction of host resistance mechanism or derived from viral priming of host metabolic pathways. The new ability to define distinct “infection states” on a pseudo-temporal manner can potentially provide valuable information regarding the dynamics of active viral infection in “real time” also in the natural environment. Clustering of individual cells based on their specific transcriptomic signatures will uncover the relationship between host metabolic states and specific phenotypes associated with differential levels of viral infection or modes of resistance in natural populations. *In situ* quantification of the fraction of infected cells, their infection and metabolic states and the fraction of resistant cells will provide important insights into the infection dynamics and may provide fundamental understating of host-virus co-existence strategies in the ocean. Resolving host-virus interaction on a single cell will provide novel sensitive biomarkers to assess the ecological impact of marine viruses and their role in regulating the fate of algal blooms in the ocean.

## Methods

### Culture growth and viral infection dynamics

Cells of the non-calcified CCMP2090 and calcifying RCC1216 *E. huxleyi* strains were cultured in K/2 medium [72] and incubated at 18°C with a 16:8 h light–-dark illumination cycle. A light intensity of 100 μM photons·m^−2^·s^−1^ was provided by cool white LED lights. Experiments were performed with exponential phase (5·10^−5^-1·106 cells·ml^−1^) or stationary phase (5·10^6^ cells·ml^−1^) cultures. *E. huxleyi* virus EhV201 (lytic) used for this study was isolated originally in [12]. In CCMP2090 experiments, *E. huxleyi* was infected with a 1:50 volumetric ratio of viral lysate to culture (multiplicity of infection (MOI) of about 1:1 infectious viral particles per cell). In RCC1216 experiments, *E. huxleyi* was infected with a 1:1000 volumetric ratio of viral lysate to culture (MOI of about 1:0.2 infectious viral particles per cell). For single-cell analysis, *E. huxleyi* cells were concentrated to 2.5·10^6^ cells·ml^−1^ by gentle centrifugation (3000 RPM, 3 min) prior to single-cell isolation. To compare between viral infection in exponential and stationary phases, stationary phase cells were diluted to similar concentration of exponential phases cells using stationary conditioned medium (5·105-1·10^6^ cells·ml^−1^) and then infected by EhV. The growth dynamics of *E*. *huxleyi* CCMP2090 strain and RCC1216 strain clones were monitored in seawater-based K/2 medium in control conditions and in the presence of the lytic viral strain EhV201. Resistant single cells were isolated after infection by mouth-pippetting over multiple passages through fresh medium under an inverted microscope. Single isolates were maintained in K/2 medium.

### Enumeration of cell and virus abundance

Cells were monitored and quantified using a Multisizer 4 Coulter counter (Beckman Coulter, Nyon, Switzerland). For extracellular viral production, samples were filtered using 0.45 μM PVDF filters (Millex-HV, Millipore). Filtrate was fixed with a final concentration of 0.5% glutaraldehyde for 30 min at 4ºC, then plunged into liquid nitrogen, and stored at −80ºC until analysis. After thawing, 2:75 ratio of fixed sample was stained with SYBER gold (Invitrogen) prepared in Tris–EDTA buffer as instructed by the manufacturer (5 μl SYBER gold in 50 mL Tris–EDTA), then incubated for 20 min at 80ºC and cooled down to room temperature. Flow cytometric analysis was performed with excitation at 488 nm and emission at 525 nm.

### Single-Cell Quantitative RT-PCR

Single cells were captured on a C1 STA microfluidic array (5–10 μm cells) using the Fluidigm C1 and imaged on IX71S1F-3-5 motorized inverted Olympus microscope (Tokyo, Japan) to examine chlorophyll autofluorescence (ex:500/20 nm, em:650 nm LP). Only wells that exhibited chlorophyll autofluorescence signal emitted from single cells were further analyzed. External RNA Controls Consortium (ERCC) spikes were added to each well in a final dilution of 1:40,000. Cells were lysed and pre-amplified cDNA was generated from each cell using the Single Cells-to-CT Kit (Life Technologies). Pooled qPCR primers and Fluidigm STA reagents were added according to manufacturer’s recommendations. Preamplified cDNA was then used for high-throughput qPCR measurement of each amplicon using a BioMark HD system. Briefly, a 2.7 μl aliquot of each amplified cDNA was mixed with 3 μl of 2X SsoFast EvaGreen Supermix with Low ROX (Bio-Rad) and 0.3 μl of 20X DNA Binding Dye Sample Loading Reagent (Fluidigm), and 5 μl of each sample mix was then pipetted into one sample inlet in a 96.96 Dynamic Array IFC chip (Fluidigm). Individual qPCR primer pairs (50 μM, Supplemental Table 1) in a 1.08 μl volume were mixed with 3 μl Assay Loading Reagent (Fluidigm) and 1.92 μl Low TE, and 5 μl of each mix was pipetted into one assay inlet in the same Dynamic Array IFC chip. Subsequent sample/assay loading was performed with an IFC Controller HX (Fluidigm) and qPCR was performed on the BioMark HD real-time PCR reader (Fluidigm) following manufacturer’s instructions using standard fast cycling conditions and melt-curve analysis, generating an amplification curve for each gene of interest in each sample (cell). Data was analyzed using Real-time PCR Analysis software (Fluidugm) with the following settings: 0.65 curve quality threshold, linear derivative baseline correction, automatic thresholding by assay (gene), and manual melt curve exclusion. Cycle threshold (Ct) values for each reaction were exported. As seen in other applications of this technology*[62]*, the data had a bimodal distribution with some cells ranging from 2.5 Ct to 30 Ct, and another set of cells with Ct >40. Similar bimodal distribution was also observed for the ERCC spikes. Accordingly, we set the minimal threshold level of detection to 30 Ct and calculated expression threshold values (Et) by linear transformation of the data so that minimal Et was zero (30 Ct). For heat map visualization, expression data was normalized by subtracting the mean of each gene and dividing it with its standard deviation across cells. Single-cell PCR data was analyzed and displayed using MATLAB (MathWorks). Additional statistical analyses were performed using The SingleCellAssay R package [62]. Calculation of number of spike molecule per Fluidigm C1 well was performed according to [61].

## Acknowledgments

We wish thank Dr. Daniella Schatz and Guy Schleyer from the Vardi lab, Dr. Roi Avraham from the Department of Biological Regulation at the Weizmann Institute of Science and Dr. Noam Stern-Ginossar from the Department of Molecular Genetics at the Weizmann Institute of Science for critical comments on the manuscript and fruitful discussion. We would also like to thank Tal Bigdary from the Design, Photography and Printing Branch at the Weizmann Institute of Science for assistance in designing the graphs for this manuscript.

## Author contributions

S.R. and A.V. conceived and designed the experiments and wrote the manuscript. S.R., M.J.F and D.P conducted the single-cell experiments. R.R. preformed single-cell statistical analysis. S.R analyzed the single-cell expression data.

